# Solution structure of the Z0 domain from transcription repressor BCL11A sheds light on the sequence properties of protein-binding zinc-fingers

**DOI:** 10.1101/2024.12.10.627839

**Authors:** Rilee E. Harris, Richard D. Whitehead, Andrei T. Alexandrescu

## Abstract

The transcription repressor BCL11A, which governs the switch from fetal to adult hemoglobin during development, is the target of the first FDA-approved CRISPR/Cas9 gene-editing therapy in humans. By targeting BCL11A, fetal hemoglobin expression is de-repressed to substitute for defective adult hemoglobin in inherited diseases including beta-thalassemia and sickle-cell anemia. BCL11A has six CCHH-type zinc-fingers of which domains 4-6 are necessary and sufficient for dsDNA binding. Here, we focus on the CCHC-type ZNF at the N-terminus of BCL11A (residues 46-72), Z0, thought to modulate oligomerization of the transcription repressor. Using NMR and CD spectroscopy, Z0 is shown to be a thermostable CCHC zinc-finger with a pM dissociation constant for zinc. The NMR structure of Z0 has a prototypical beta-beta-alpha fold, with a hydrophobic knob comprising about half the structure. The unusual proportion of hydrophobic residues in Z0 led us to investigate if this is more general in zinc-fingers that do not bind dsDNA. We used the ZF and WebLogo servers to examine sequences of zinc fingers with demonstrated DNA-binding function, non-binders, and the CCHC-type family of protein-binders. DNA-binders are distinguished by contiguous stretches of high-scoring zinc-fingers. Non-DNA-binders show a depletion of polar residues at the positions expected to contact nucleotides and increased divergence from sequence consensus making these domains more likely to be annotated as atypical, degenerate, or to be missed as zinc-fingers. We anticipate these sequence patterns will help distinguish DNA-binders from non-binders, an open problem in the functional understanding of zinc-finger motifs.

## 1. INTRODUCTION

Several inherited blood diseases, including β-thalassemia and sickle cell anemia occur due to mutational defects in the β-chain of hemoglobin (Hb). Adult Hb is a protein comprised of two α and two βchains arranged in a tetrameric α_2_β_2_ quaternary structure (Perutz 1964; Weatherall *et al*. 2006). The oligomeric composition of Hb changes during normal human development (Sankaran and Orkin 2013). In embryonic stages, Hb uses α-like ζ-globin and β-like ε-globin chains to form a ζ_2_χ_2_ tetramer (Wilber *et al*. 2011). By twelve weeks of gestation almost all Hb is composed of α-chains and β-like γ-chains to form α_2_γ_2_ fetal hemoglobin (HbF) optimized for O_2_ transport from the mother to the fetus. The switch between fetal and adult Hb (HbA) starts in the last trimester, with adult α_2_β_2_ Hb becoming the dominant form of Hb after birth (Sankaran and Orkin 2013). The switch between HbF and HbA is governed by the transcriptional repressor B-cell lymphoma/leukemia 11A (*BCL11A*), a zinc-finger (ZNF) transcription factor (TF) that represses the expression of *γ-globin* (and therefore HbF) by binding directly to the *γ-globin* gene promoter elements (Liu N *et al*. 2018; Sankaran and Orkin 2013; Shen *et al*. 2021; Zhang *et al*. 2024). In addition to HbF silencing, *BCL11A* has roles in brain and hematopoietic system development, and may contribute to human diseases such as type II diabetes and several types of cancers (Katnik *et al*. 2023; Satterwhite *et al*. 2001; Seigfried and Britsch 2024; Sunami *et al*. 2022; Yin J *et al*. 2019; Zhou *et al*. 2020). Inherited intellectual development disorders are associated with mutations in *BCL11A* (Peron *et al*. 2024; Shen *et al*. 2021), including the missense mutations C48F, T47P, M53P, and H66Q in the first zinc finger, Z0. Carriers of these mutations often display HbF persistence as a biomarker (Peron *et al*. 2024). *BCL11A* was formerly listed by NIH/NCATS as an understudied rare disease protein under the program RFA-TR-22-030.

There has been considerable recent interest in *BCL11A* as a therapeutic target. Fetal HbF can functionally substitute for adult HbA (Frangoul *et al*. 2024; Sankaran and Orkin 2013; Song *et al*. 2024). Thus, abrogation of *BCL11A*, the transcriptional repressor of *γ-globin* gene expression, leads to increased levels of the beneficial fetal hemoglobin molecule (α_2_γ_2_) that can substitute for adult (α_2_β_2_) hemoglobin in inherited disorders like sickle cell disease and β-thalassemia (Zheng and Orkin 2024). Such an approach forms the basis of a joint venture between the pharmaceutical company Vertex and Harvard University that in 2023 that led to Casgevy, the first FDA-approved CRISPR/Cas9 gene-editing therapy in humans (Frangoul *et al*. 2024; Sheridan 2024). The Casgevy therapy is an exagamglogene autotemcel treatment for sickle cell disease and β-thalassemia in which the patient’s blood stem cells are genetically edited and returned in a single infusion after treatment with high-dose chemotherapy to remove bone marrow cells (Demirci *et al*. 2021; Frangoul *et al*. 2024). The CRISPR/Cas9 genetic modification does not repair the *β-globin* gene but disrupts expression of *BCL11A* (Demirci *et al*. 2021; Frangoul *et al*. 2024; Fu *et al*. 2022; Song *et al*. 2024), allowing increased expression of the HbF γ*-globin* gene that can functionally substitute for defective *β-globin* chains in adult HbA. While the results of Casgevy have been positive, the search continues for methods to control *BCL11A* function that circumvent the need for gene editing (Zheng and Orkin 2024). These alternative approaches include miRNA silencing (Esrick *et al*. 2021), proteolysis targeting chimera (PROTAC) strategies to target the BCL11A protein for proteosome degradation via ubiquitination (Mehta *et al*. 2022), evolved nanobodies directed to the transcription repressor (Yin M *et al*. 2023), as well as more conventional destabilization of BCL11A by small molecule drugs (Shen *et al*. 2021; Yang JP *et al*. 2024; Zheng and Orkin 2024).

Recent studies have started to reveal key information on the structure and biophysics of *BCL11A*. An X-ray crystallographic study looked at a complex of the last three of six CCHH ZNF domains, ZNF4-6, in complex with a 12 bp DNA duplex corresponding to the proximal promoter region of the *γ-globin* gene that is repressed by BCL11A. The work delineated key interactions used by the ZNF domains to recognize the cognate DNA sequence (Yang Y *et al*. 2019). These interactions were further verified using structure-function analyses of individual ZNF domain disruption using CRISPR-Cas9 in adult erythroid cells (Rajendiran *et al*. 2024). The ZNF4-6 fragment was characterized in its free and DNA-bound forms using NMR, X-ray crystallography, and MD simulations (Viennet *et al*. 2024). An X-ray structure of ZNF6 in complex with a nanobody engineered to target *BCL11A* for degradation was recently determined (Yin M *et al*. 2023).

In addition to six CCHH-type ZNFs in *BCL11A*, of which ZNF4-6 are necessary and sufficient for DNA binding (Liu N *et al*. 2018; Shen *et al*. 2021; Yang Y *et al*. 2019), a putative CCHC-type ZNF was described at the N-terminus of the transcription factor (Grabarczyk *et al*. 2018; Shen *et al*. 2021) that we will henceforth call Z0. The Z0 domain is thought to function in the homodimerization of *BCL11A*, which plays a part in controlling its transcription regulation activity (Grabarczyk *et al*. 2018; Liu H *et al*. 2006; Susemihl *et al*. 2021). Due to marked sequence differences from the ZNF consensus the UniProt database (The_UniProt_Consortium 2023) does not recognize the Z0 domain as a ZNF (residues 46-72 in UniProt entry Q9H165). Similarly, the ZNF-motif prediction server (https://zf.princeton.edu/logoMain.php) (Persikov and Singh 2014) is unable to detect a ZNF, scoring the 46-72 sequence segment with a zero. Given the importance of Z0 in the dimerization function of BCL11A (Grabarczyk *et al*. 2018) and the potential of the domain as a drug target (Shen *et al*. 2021), we present in this report a biophysical and structural characterization of the domain. Using NMR and CD spectroscopy, we show that the domain folds in the presence of Zn(II) and is a genuine thermostable ZNF. UV-Vis spectrophotometry of the Z0-Co(II) complex is used to show that the domain is a CCHC-type ZNF that binds metal with a tetrahedral coordination sphere. We determine the K_d_ for Zn(II)-binding and obtain NMR assignments and a solution structure for the Z0 domain that could be useful in future drug-targeting of the domain. Finally, existing sequence analysis programs (Crooks *et al*. 2004; Persikov and Singh 2014) are used to explore the open question of what sequence and structural properties distinguish whether ZNFs function as nucleic acid-binding domains such as the *BCL11A* ZNF4-6, or protein binders such as the Z0 domain.

## 2. RESULTS AND DISCUSSION

### 2.1 Zn^2+^-binding through a CCHC ligating site induces folded structure in the Z0 domain

We employed a variety of spectroscopic methods to ascertain the Zn^2+^ binding properties of a L46-N72 fragment of BCL11A corresponding to the Z0 domain (Fig. 1A). Figure 1B shows circular dichroism (CD) spectra of Z0 in the presence and absence of ZnSO_4_. When Zn^2+^-bound, the peptide has a CD spectrum typical of a folded polypeptide with nearly equal minima at 208 and 224 nm and a maximum at 192 nm. In the absence of metal, the CD spectrum shows a deep minimum at 201 nm typical of an unfolded peptide but also a second minimum at 224 nm suggestive of residual α-helical structure. At pH 2 in the absence of Zn^2+^ the CD spectrum more closely approaches that of a random coil, showing a deeper 201 nm minimum at the expense of the 224 minimum. The analysis indicates that the Z0 peptide folds in the presence of Zn^2+^, but that there is a small amount of residual α-helix structure in the absence of metal at neutral pH, that becomes unfolded at acidic pH.

**Figure 1.**
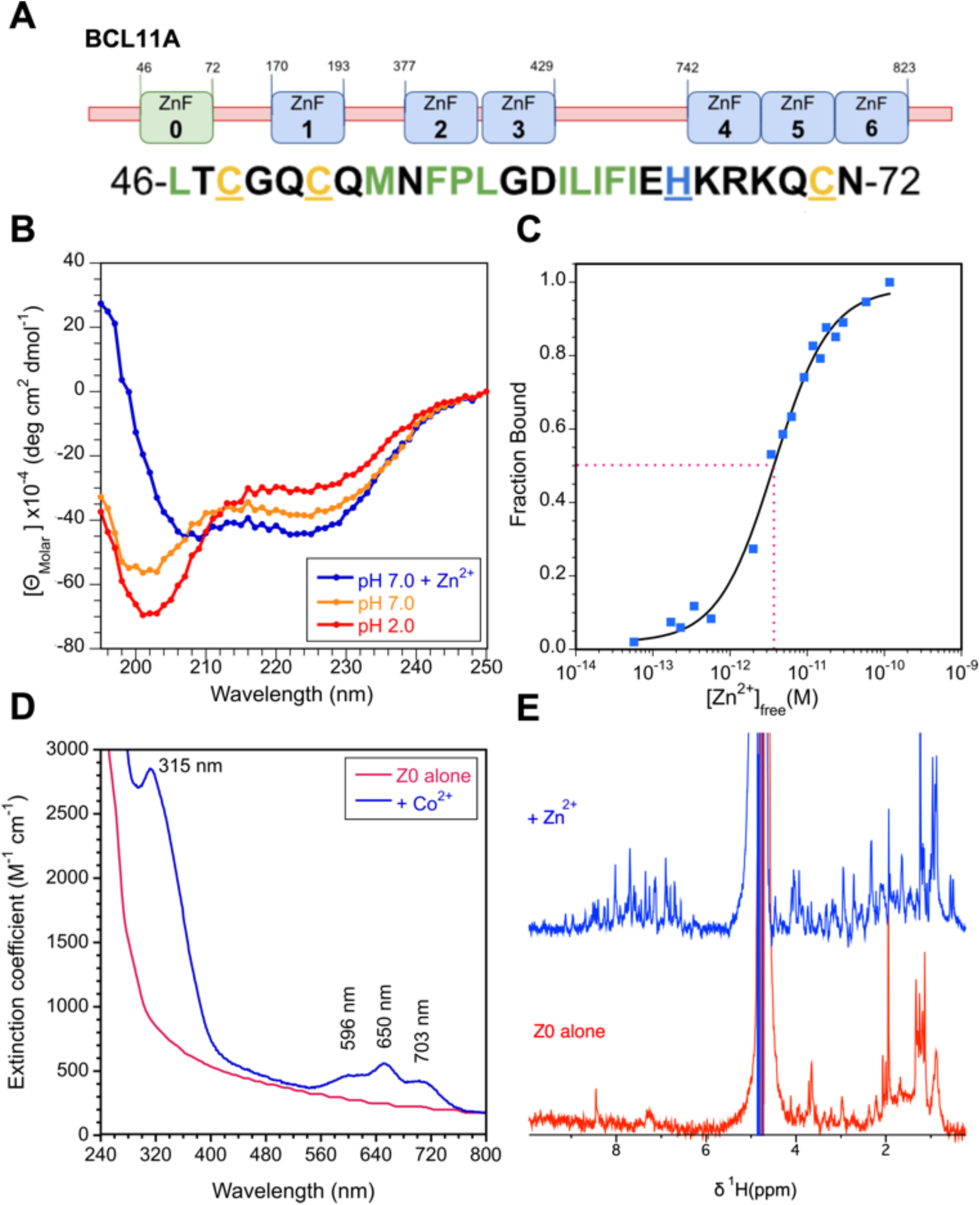
Metal binding of Z0. (A) Domain organization of BCL11A and sequence of the Z0 domain peptide fragment studied in this work. In the sequence metal ligands are underlined, and hydrophobic residues are indicated in green type. (B) CD spectra of Z0 with Zn^2+^ and in the absence of the metal at neutral and acidic pH. (C) Zn^2+^ titration in competition with the chelator EGTA at pH 7.0 and a temperature of 20 °C used to determine the *K*_d_ value of (3.7 ± 0.3) x 10^-12^ M for metal binding (dotted red line). (D) UV-Vis spectrum of Z0 without and with CoCl_2_ used to obtain information on metal coordination at a temperature of 25 °C. (E) 1D ^1^H NMR spectra of Z0 in the absence and presence of equimolar ZnSO_4_ at pH 6.8, 25 °C.

Using the difference in CD ellipticity at 200 nm between the apo and holo peptide at pH 7.0, we used a previously published competition method using the metal chelator EGTA to determine a *K*_d_ value of (3.7 ± 0.3) x 10^-12^ M for Zn^2+^ binding to the Z0 domain (Fig. 1C). The thermal stability of the Zn^2+^-bound state of Z0 was characterized using a temperature titration between 20 and 90 °C at pH 7.0. With increasing temperature, the CD spectra showed only small gradual changes typical of a pre-unfolding transition indicating that the Z0 domain is highly thermostable (Supplementary Figure 1). Cellular thermal shift assays (CETSA) found a melting temperature of 49.7 °C for the BCL11A XL isoform *in vivo* (Shen *et al*. 2021), a value more than 40 degrees lower than the stability of the isolated Z0 domain measured *in vitro* (Fig. S1). These observations suggest targeting drugs to destabilize the Z0 domain might be more challenging than disrupting intermolecular interactions involving Z0.

UV-vis spectrophotometry of the Co^2+^-bound Z0 fragment was employed to obtain information on the metal coordination site (Fig. 1D). The spectrum in the presence of Co^2+^ has a ligand-to-metal charge transfer (LMCT) transition with an ε_315_ of 2842 M^-1^ cm^-1^. Each S^-^-Co^2+^ bond is expected to contribute 900-1200 M^-1^ cm^-1^ to the LMCT extinction coefficient (Nomura and Sugiura 2002), so the value for Z0 is consistent with three cysteines coordinating Co^2+^. The d-d orbital transition region in the low energy region of the UV-Vis spectrum has three bands at 596, 650, and 703 nm. The positions of these bands imply a Cys3His ligand set for the metal (Sivo *et al*. 2017), while their extinction coefficients greater than 400 M^-1^ cm^-1^ are consistent with a tetrahedral coordination sphere (Nomura and Sugiura 2002). Taken together, UV-Vis spectroscopy of the Co^2+^-Z0 complex indicates the ZNF binds metal through its CCHC residues in a tetrahedral coordination geometry.

Figure 1E compares 1D 1H NMR spectra of Z0 at pH 6.8 with and without equimolar ZnSO_4_. In the presence of Zn^2+^ there is an increase in chemical shift dispersion indicating the peptide adopts a folded structure upon metal binding. Backbone amide proton chemical shifts range from 7.61 ppm for D14 to 9.15 ppm for C3, and the methyl peaks groups of L1 resonate at 0.51 and 0.56 ppm due to the ring-current shift from F18. In the presence of Zn^2+^, more amide protons are observed between 7 and 9 ppm due to increased solvent exchange protection in the folded structure.

### 2.2 The NMR structure of the Z0 domain has a hydrophobic knob

NMR structure determination of Z0 proved challenging because of the low solubility of the Zn^2+^-bound peptide, as well as time-dependent degradation of the NMR spectrum due to aggregation at high concentrations above 0.5 mM. Under acidic conditions below pH 4.5 the Z0 peptide is soluble in water, but in the pH range 5.5 to 7.0 typically used for NMR spectroscopy the peptide often precipitated in the presence of equimolar ZnSO_4_. Despite precipitation, it was still possible to obtain NMR spectra. The concentration of soluble peptide remaining in the supernatant after removing the precipitate by sedimentation was determined to be 40 µM using a bicinchoninic acid (BCA) assay (Walker 2002). We therefore did most of our subsequent NMR studies on 40 µM concentrations of the Zn^2+^-bound peptide to avoid aggregation. Under these conditions Zn^2+^-bound Z0 is soluble and stable for NMR runs of at least 1-2 days. The low 40 µM concentration, however, posed challenges for both NMR sensitivity and suppression of the H_2_O solvent signal, since our peptide samples were not isotopically enriched. 2D TOCSY and NOESY experiments in H_2_O enabled us to obtain sequential assignments for Zn^2+^-bound Z0, but the NOESY spectrum contained relatively few structurally useful distance restraints (5 long-range and 10 medium-range NOEs). By contrast, the problems associated with solvent suppression at low peptide concentrations were circumvented for NOESY spectra of Zn^2+^-bound Z0 in D_2_O. We were able to obtain a larger number of structurally relevant distance restraints in NOESY spectra of Z0-Zn^2+^ dissolved in 99.96% D_2_O (26 long-range and 22 medium-range NOEs). The downfield region of the NOESY spectrum in D_2_O is shown in Fig. 2A, highlighting the quality of the data and NOE assignments for aromatic and Hα protons. Figure 2B summarizes the chemical shift and short-range NOE data that were used to calculate a consensus secondary structure for Zn^2+^-bound Z0.

**Figure 2.**
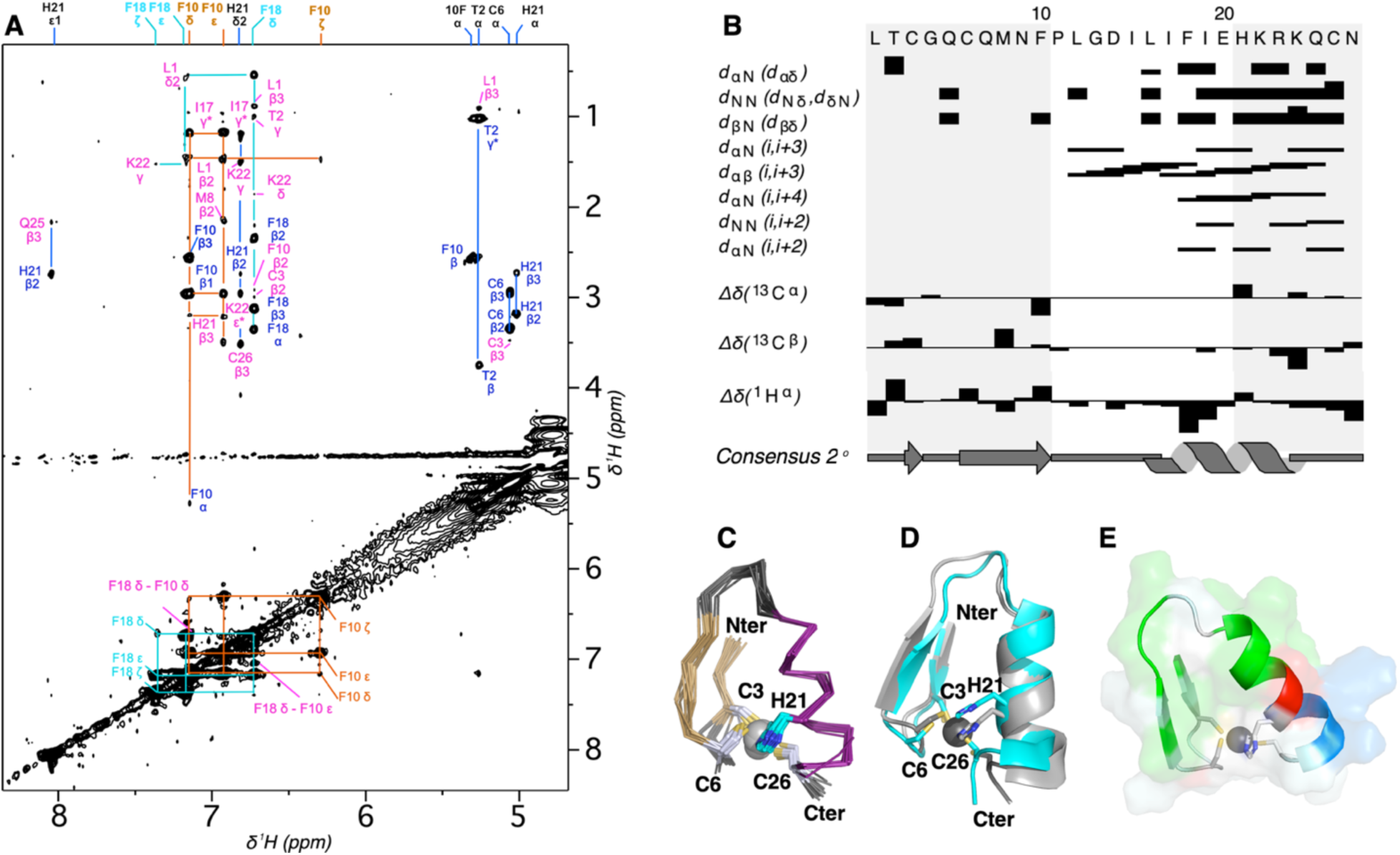
NMR structure of Zn^2+^-bound Z0. **(A)** Aromatic region of a 200-ms mixing time NOESY spectrum recorded on a sample in D_2_O showing intra-(blue) and inter-residue (pink) NOE distance contact assignments. **(B)** Wüthrich diagram calculated with the program DANGLE (Cheung, *et al*. 2010) summarizing chemical shifts indices and short-range NOEs pertaining to secondary structure. (C) Ensemble of 20 lowest energy NMR structures showing the Z0 main-chain and Zn^2+^-ligating residues. (D) best fit superposition of the Z0 NMR structure closest to average (gray) and the Colabfold predicted structure (cyan). The all-atom RMSD between the two structures is 2.11 Å (backbone C(,C,N,O is 1.60 Å). (E) Surface model of the Z0 structure with the hydrophobic knob (residues L1, M8, 10-FPL-12, and 15-ILIFI-19) shown in green. The only charged residues in Z0 are at the C-terminus: E20 and 22-KRK-24 shown in red and blue, respectively.

Despite the technical challenges of working at low peptide concentrations, the NMR structures of Z0-Zn^2+^ readily converged to a unique solution, with no irreconcilable restraint violations. Figure 2C shows the top 20 lowest-energy NMR structures for Z0. The secondary structure is well-defined and consists of a β-hairpin (gold) and α-helix (purple) in the folding arrangement typical of ZNFs. The four Zn^2+^ ion ligands C3, C6, H21, and C26 are also well-defined in the structure. The experimental NMR structure agrees with that predicted by ColabFold (Mirdita *et al*. 2022), giving RMSDs of 1.3 Å and 2.1 Å for backbone and all-atoms, respectively (Fig 2D, Table S1). In light of the NMR structure, we can examine the mutations associated with inherited intellectual disability that occur in this domain (Peron *et al*. 2024; Shen *et al*. 2021). The T47P and M53P mutations introduce proline ‘structure breakers’ in strands β1 and β2, respectively. The C48F and H66Q mutations remove the first and third ligands for Zn^2+^ respectively (Fig. 1A). We caution that loss of one of the four ligands can sometimes be tolerated and does not necessarily imply an inability of ZNFs to bind Zn^2+^ or fold (Rua and Alexandrescu 2024).

We compared the Z0 NMR structure to that of the CCHC-type ZNF from the protein ZNF750 that we recently determined (Rua *et al*. 2023), and obtained a backbone RMSD of 2.1 Å. Even though the Z0 domain and the one from ZNF750 are both CCHC-type ZNFs, the Z0 zinc finger has significant deviations from the amino acid sequence consensus for this fold. In the Z0 domain (Fig. 1A), (i) the conserved aromatic two positions upstream of the first cysteine is missing, (ii) the conserved hydrophobic residue three positions before the histidine is replaced by an aromatic phenylalanine, and most importantly (iii) the ‘finger’ region between the second and third Zn^2+^ ligands is 14 rather than the typical 12 residues. The extra two residues are accommodated by an extra bulge in the loop between the β-hairpin and α-helix components of the ZNF. These features combine to make Z0 unrecognizable as a ZNF with either the UniProt database (accession code Q9H165 residues 46-72), or the ZF-prediction server (Persikov and Singh 2014). Despite the deviations from the sequence consensus the Z0 domain adopts a structure similar to the CCHC ZNF from ZNF750, implying a malleable folding motif that can accommodate considerable sequence diversity.

Another unusual feature of the Z0 motif is the presence of a hydrophobic knob that comprises about half of the structure (green in Fig. 2E). This hydrophobic region is likely to participate in the oligomerization function of the Z0 domain (Grabarczyk *et al*. 2018). The hydrophobic portion of the molecules is comprised of residues L1, M8, 10-FPL-12 and a five amino acid stretch 15-ILIFI-19 at the start of the α-helix portion of the structure. By contrast the C-terminus of the α-helix has a string of six polar residues 20-EHKRKQ-25, with H21 the sole histidine ligand for Zn^2+^.

We looked for NOE distance contacts in our NOESY data that could provide clues about the structure of a dimeric version of Zn^2+^-bound Z0. Out of 174 assigned NOEs only four were consistently violated in initial monomer structures: F18 Hδ - M8 Hβ, I15 NH - P11 Hγ, H21 Hδ2 – M8 Hα, and H21 Hδ2 – I15 Hγ1. Of these, the last three have alternative possibilities consistent with short distance contacts in the Z0-Zn^2+^ monomer structure: I15 NH - I17 Hβ, F18 Hε - I15 Hα, and F18 Hε - I19 Hγ1. Moreover, H21 is highly unlikely to be involved in dimer contacts since it is ligating zinc. Only the F18 Hδ - M8 Hβ lacked alternative assignments, however, the corresponding distance of 6.5 Å could be plausible if the NOE has contributions from spin diffusion or aromatic ring flipping of F18. Because of the ambiguities associated with the four violated NOEs we excluded them from structure calculations, but none were consistent with the concomitant presence of an oligomer in the sample. The remaining 170 NOEs were all compatible with the monomer NMR structure shown in Fig. 2C, that agrees with the structure predicted by ColabFold (Fig. 2C). If there are Z0 oligomers present at the 40 µM sample concentration used for our studies, the amounts are too low to contribute to the NOESY spectrum.

We then examined whether selective NMR line-broadening at higher Z0 concentrations could give structural information on dimerization. For this experiment we prepared a 40 µM Z0 sample with 40 µM ZnSO_4_ and divided it into various aliquots that after lyophilization and dissolution in D_2_O would give 0.04, 0.71, 1.1, and 2.1-mM concentrations of the Zn^2+^-bound Z0 domain. At Z0 concentrations greater than 0.7 mM we observed line broadening throughout the 1D ^1^H NMR spectra (Fig. S2), suggesting the oligomerization interface involves contacts distributed throughout the molecule rather than limited to only a subset of residues.

Finally, we tried to model the structure of dimeric Z0-Zn^2+^ with the AlphaFold3 server (https://alphafoldserver.com) (Abramson *et al*. 2024). Of the five models predicted by Alphafold3, three (M2-M4) are structurally convergent with an average RMSD of 1.3 Å (Fig. S3A). The top ranked model M1, however, gives an average RMSD of 10 Å to models M2-M4. In all five models, the hydrophobic region 15-ILIFI-19 at the N-terminus of the α-helix is part of the dimerization interface. Compared to models M2-M4, model M1 gives a more extensive dimerization interface involving the α-helix as well as hydrophobic segments from the β-hairpin. By contrast, models M2-M4 seem less energetically plausible as some of the hydrophobic sidechains are facing solvent (Fig. S3B). Model M5 has the lowest AlphaFold pLDDT confidence and gives an average RMSD of about 5 Å to the other models. Thus, the predicted AlphaFold3 models did not converge to a unique solution. We next tried the same approach with oligomerization states ranging from trimers (n=3) to nonamers (n=9). Each of the putative oligomeric structures showed similar heterogeneity amongst the top 5-scored AlphaFold3 structures, with at least one of the top five scorers representing an outlier with an RMSD greater than 5 Å to the others. Perhaps the structural heterogeneity in the AlphaFold3 predictions suggests oligomerization plasticity, in as much as Z0 can adopt a number of oligomerization states (Grabarczyk *et al*. 2018; Zheng *et al*. 2024) including the monomer studied in this work.

### 2.3 Non-DNA-binding ZNFs have more varied sequence preferences than DNA-binding counterparts

It is becoming increasingly appreciated that the types of ZNFs usually associated with dsDNA-binding functions in TFs can also have protein binding functions (Brayer and Segal 2008; Iuchi 2001; Laity *et al*. 2001; Ni *et al*. 2020; Persikov and Singh 2011; Schafer *et al*. 1994). In several TF proteins, ZNFs are modularly arranged in arrays, with some examples having 30 or more copies. Yet only a small subset of the ZNFs participate in dsDNA binding, with the remainder having unknown, protein-binding, or structural roles (Iuchi 2001; Ni *et al*. 2020). Here, we consider that ‘structural roles’ could describe protein-interaction within a TF rather than interactions between two or more proteins. The sequence or other features that specify dsDNA versus non-DNA-binding functions are unknown.

The presence of a hydrophobic knob in the Z0 structure led us to examine if DNA- and non-DNA-binding ZNFs might have different amino acid sequence signatures. To distinguish between DNA and non-DNA binding ZNF domains we constructed four datasets. The first dataset, *DNA-binders X-ray*, had arrays of two or more ZNFs in complex with dsDNA from X-ray structures in the Protein Data Bank (PDB). In total, this dataset included 102 CCHH-type ZNF domains bound to DNA, from 26 separate human TF X-ray structures (Table S2). The remainder of the ZNF domains catalogued in the UniProt entries of these proteins were not assigned as non-DNA binders, since the lack of a crystal structure does not exclude the possibility that a ZNF could bind DNA but fail to crystalize.

The next two databases *DNA-binders Δ* and *Non-DNA Δ* were constructed from 18 literature papers reporting on mutagenic deletion analysis to distinguish ZNF domains necessary and sufficient for dsDNA-binding from ZNFs not implicated in DNA binding. Each of the 18 TFs had between 4 and 30 ZNFs for a total of 148. Of these, 57 were DNA-binders and 91 were non-DNA-binders (Table S3). Since the ZNFs arrays in these TFs likely arose through gene duplication, they offer a stringent test for distinguishing DNA-binders from non-binders based on amino acid sequences.

The last dataset, *CCHC protein-binders*, included 28 CCHC-type ZNFs for which there is literature evidence for protein rather than DNA-binding functions (Table S4). In previous work we referred to this ZNF subfamily as ‘CCHC-finger’ (Rua *et al*. 2023) to distinguish it from the shorter ‘knuckle-CCHC’ subfamily that has a C-X2-X4-H-X4-C sequence spacing and typically binds ssRNA and ssDNA. The CCHC-finger subfamily (Ω)-X-C-X_2-5_-C-X_3_-ψ-X_5_-λφϕ-X_2_-H-X_3-5_-H, where Ω and ψ are aromatic and hydrophobic residues, respectively, has the same sequence spacing as found for the CCHH-fingers that occur in TFs and nearly identical structures (Rua *et al*. 2023). The only substantial difference is the switch of the last Zn^2+^ ligand from a histidine to a cysteine. Early protein design experiments showed that histidine and cysteine are structurally interchangeable in the last Zn^2+^ ligand position, and that substitution of a cystine for a histidine at the last Zn^2+^-ligand position of ZNF3 from transcription repressor BKLF does not affect its ability to bind dsDNA (Simpson *et al*. 2003). Yet naturally occurring CCHC-fingers found in proteins such as Fog, MOZ, and NEMO all have protein-binding functions (Cordier *et al*. 2008; Matthews *et al*. 2000). In fact, we are unaware of any examples of a CCHC-finger with a DNA-binding function.

We first analyzed the ZNF sequences in the four datasets using the ZF-motif server (Persikov *et al*. 2009; Persikov and Singh 2014). This program predicts the DNA-binding specificities of CCHH ZNFs. As a first step of this process a Hidden Markov Model protein sequence homology search (HMMER) is performed against the CCHH-ZNF consensus defined in the protein database family (Pfam) to identify and score ZNF domains. HMMR bit scores > 17.7 constitute the most confident ZNF predictions (Persikov and Singh 2011). We used the HMMR bit score to look for differences between our four ZNF datasets, as summarized in the dot plot of Figure 3A. The difference in the means of the first two datasets, the DNA-binders from X-ray structures and those from deletion analyses, is not significant as expected since both sets of ZNFs bind DNA. Significant differences (p <0.05) are observed for all other pairwise comparisons of the means. The group *Non-DNA Δ* have a mean score of 21.7, approaching the 17.7 cutoff threshold for ZNF identification by the ZF-server. The *CCHC protein-binders* group gives a mean score of 9.4 that is below the threshold for confident ZNF identification. Since the HMMR bit score essentially tries to match sequences to the aforementioned idealized consensus CCHH ZNF sequence, these observations indicate that the sequences of both non-DNA binding subtypes have considerable differences form DNA-binding ZNFs.

**Figure 3.**
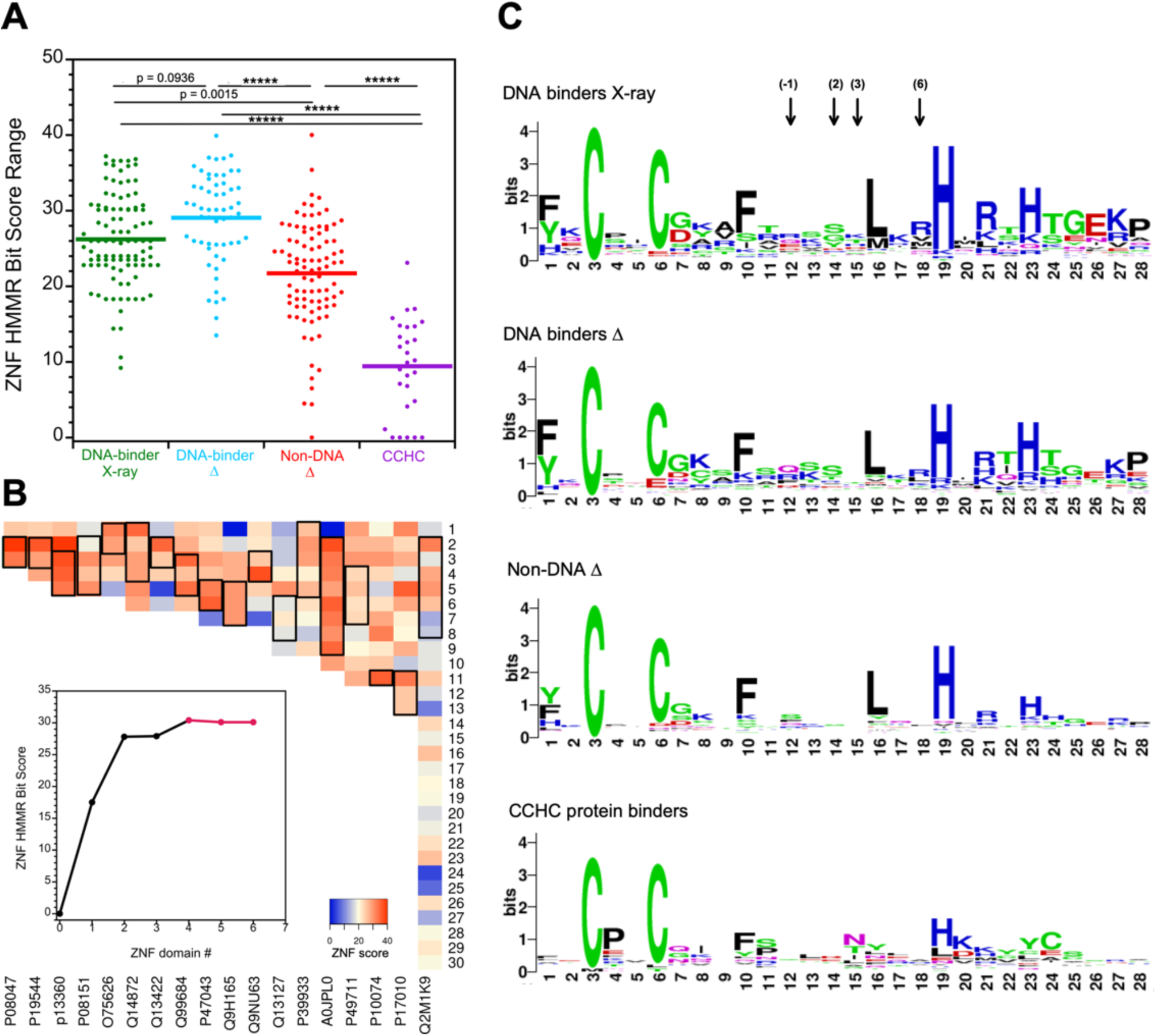
Sequence properties of DNA- and protein-binding CCH(H/C) ZNFs. **(A)** Dot plots of ZNF HMMR scores calculated with the ZF server. Each point represents a ZNF domain from one of the four indicated datasets: DNA-binders from X-ray structures (mean score = 26.2 ± 5.9, n = 102), DNA-binders from deletion analyses (mean score = 29.1 ± 6.0, n = 57), non-DNA-binders from deletion analyses (mean score = 21.7 ± 6.9, n = 91), CCHC protein-binders (mean score = 9.4 ± 6.3, n = 28). The statistical significance of the difference between means calculated with an Anova / post-hoc Tukey HSD test is given by the p-values, with ***** indicating p < 0.00001. **(B)** Heat maps summarizing ZNF HMMR scores of individual domains from the literature database of 18 TFs where mutational deletion analysis was used to identify DNA binding ZNFs (indicated by black rectangles). The TFs are identified by their UniProt accession ID at the bottom of the figure and are listed in order from the smallest (3) to largest (30) number of ZNF domains. The inset graph shows BCL11A. The first ZNF is Z0, the subject of the current study, and Z4-Z6 (red) are the domains required for dsDNA binding. **(C)** Amino acid logos for each of the indicated ZNF datasets. The arrows in the top logo indicate the positions of sites that contact nucleotides in DNA-binding ZNFs, numbered from the start of the ZNF α-helix (-1, 2, 3, 6).

To gain further insight, the ZNF scores for the 18 TFs analyzed by deletion mutagenesis to distinguish DNA-binders from non-binders were visualized using a heatmap approach (Fig. 3B). Each TF corresponding to an individual column in Fig. 3B, is sorted from the smallest number (3) to the largest number (30) of ZNFs. Arranged this way, it is apparent that the TFs with a smaller number of ZNFs (redder) have larger average scores than those with a greater number of ZNF domains. In TFs with a larger arrays of ZNFs, the domains with the higher scores corresponding to stronger DNA-binding motif matches appear to be diluted by ZNFs with more divergent sequences. The black boxes in the heatmap of Fig. 3B enclose the domains determined to bind DNA in the literature deletion analyses. Not only do the DNA-binding domains typically give higher scores (redder), but the heatmap analysis shows that they tend to occur in clusters of high-scoring ZNFs. The smallest of these clusters have 2-3 contiguous ZNF domains, which likely correspond to a minimal dsDNA-binding unit (Persikov and Singh 2011). Single ZNFs are typically insufficient to bind dsDNA with specificity (Klug 2010; Lee *et al*. 1991; Omichinski *et al*. 1997). The clustering of dsDNA-binding ZNFs could satisfy structure-function requirements in TFs. Non-contiguous ZNF clusters would give a discontinuous dsDNA-binding site, that would need to position multiple ZNF clusters interrupted by a non-interacting protein segments, on the same strand of dsDNA. The segregation of scores for DNA-binding domains is illustrated with the example of BCL11A in the inset of Fig. 2B, although for several other TFs the demarcation between the DNA- and non-DNA-binding ZNF scores is less clear cut. An interesting example of why ambiguity can persist in the score demarcation is provided by the P49711 entry in Fig. 2B that represents CTCF (CCCTC-binding factor). CTCF uses different combinations of ZNFs to bind to diverged promoter sequences (Filippova *et al*. 1996). Thus, the distinction between DNA-binding and non-binding segments can be confounded when TFs are multifunctional with binding sites that depend on biological context, or when they contain evolutionary vestiges of ancestral binding sites.

We next examined what sequence features distinguished dsDNA-binding ZNF from non-binders using the WebLogo server (https://weblogo.berkeley.edu/logo.cgi) (Crooks *et al*. 2004). The sequence logos for the *DNA-binders X-ray* and *DNA-binders Δ* datasets are nearly identical since both are composed of dsDNA-binding ZNFs. These sequence logos largely correspond to the consensus sequence preference (Ω)-X-C-X_2-5_-C-X_3_-(Ω)-X_5_-ψ-X_2_-H-X_3-5_-H, originally formulated based on dsDNA-binding ZNFs from the transcription factor TFIIIA (Klug 2010; Miller *et al*. 1985).

By contrast, the sequence logos shows important differences for the last two sets of non-DNA-binders. In the ‘canonical model’ for dsDNA recognition, four residues from each ZNF reads out four nucleotides from dsDNA (Klug 2010; Persikov and Singh 2011). The ZNF residues are numbered from the start of the α-helix in the structure, as indicated at the top of Fig. 3C. Residues -1, 3, and 6 recognize a nucleotide triplet on the primary 5’ to 3’ strand of dsDNA, while residue 2 contacts a base on the complementary noncoding strand (Klug 2010; Persikov and Singh 2011). These four sites corresponding to positions 12, 14, 15, and 18 in the sequence logo diagrams of Fig. 3C are enriched in polar and charged amino acids, as expected for residues that would hydrogen-bond to polar nucleic acid bases. For the two non-DNA binding datasets there is a depletion of polar residue preferences, giving the sites 12, 14, 15, and 18, a more hydrophobic character (Fig. 3C). Even more pronounced differences occur for the *CCHC protein binder* dataset. The preferences for an aromatic residue at position 1 and a hydrophobic residue at position 16 are largely absent for the CCHC ZNFs. An apparently conserved proline at position 28 of CCHH ZNFs that may constitute a demarcation between successive ZNF domains (Laity *et al*. 2001), is depleted in the CCHC group. Two other differences of unknown significance in the *CCHC* group are a preference for a small polar asparagine or threonine at position 15, and a preference for proline at position 4 immediately following the first cystine Zn^2+^-ligand.

A final difference between the ZNFs has to do with the last two Zn^2+^-ligands which have a strong preference for positions 19 and 23 in the DNA-binders but show more variability in the non-DNA binders. We previously did a comprehensive analysis of the sequence separation of the last two Zn^2+^-ligands in ZNFs (Rua and Alexandrescu 2024), and found that while sequence intervals between three and five residues occur, the shorter three-residue separation is strongly preferred. In the *non-DNA Δ* and *CCHC* groups the preference for the last Zn^2+^-ligand at position 23 is decreased implying greater variability in the spacing of the last two Zn^2+^-ligands. Moreover, in the *CCHC* group the histidine preference for the third Zn^2+^-ligand at position 19 has also decreased, indicating greater variability in the length of the highly conserved 12-residue ‘finger’ segment between the second cysteine (position 6) and first histidine (position 19). Given the increased sequence differences from consensus for the *CCHC* and *non-DNA Δ* groups, these ZNFs are more likely to be classified as ‘atypical’, ‘degenerate’ or to not be identified as ZNFs in sequence motif searches, as is the case for the Z0 domain of BCL11A. The number of CCHC and non-DNA-binding CCHH ZNFs may thus be currently underestimated, as we previously hypothesized (Rua *et al*. 2023).

In summary the patterns we describe could prove useful in discerning DNA-binding from non-DNA-binding ZNFs, which remains an outstanding problem in the functional understanding of these protein motifs. DNA-binding ZNFs are characterized by high HMMR bit scores on the ZF-server (Persikov and Singh 2014) since the score is based on a sequence match to a consensus defined for DNA-binding ZNFs (Klug 2010). A string of high-scoring ZNFs can increase the confidence of a DNA-binding site prediction, since at least 2-3 or more contiguous ZNFs are typically required to recognize a specific DNA sequence. By contrast protein-binding ZNFs are characterized by less polar character for the residues corresponding to the canonical dsDNA binding site, and greater divergence of the sequences from consensus that include loss of conserved aromatic and hydrophobic groups, and atypical spacings between Zn^2+^-ligands.

## 3. MATERIALS AND METHODS

### 3.1 Materials

The 27-residue Z0 peptide corresponding to the fragment 46-72 of human BCL11A (UniProt Q9H165) was custom synthesized at 95% HPLC purity by Biomatik (Kitchener, Canada). Mass spectrometry of the peptide gave an observed mass consistent with the theoretical expected mass of 3,151 Da. Peptide concentrations for solution studies were determined using the BCA assay (Walker 2002). ZnSO_4_•H_2_O (purity ≥ 99.9%) and CoCl_2_•6H_2_O (purity ≥ 99%) were from Sigma (St. Louis, MO).

### 3.2 Spectrophotometric measurements of metal binding

CD experiments were performed on an Applied Photophysics Chirascan V100 Spectrometer (Surrey, UK) using a 1 mm cuvette, a 1 nm bandwidth, a 1 nm scan step size, and 5 s/point data averaging. The Z0 concentration was 40 µM in 10 mM sodium phosphate buffer, containing 0.2 mM of the reducing agent TCEP (tris(2-carboxyethyl)phosphine) to prevent disulfide formation. Biding curves to obtain a *K*_d_ for Zn^2+^ were measured at 20 °C and pH 7.0, by addition of ZnSO_4_ in a competition assay with 10 mM EGTA chelator (Ivanova *et al*. 2008; Rua and Alexandrescu 2024). The *K*_d_ value was calculated from the EGTΑ competition experiments as previously described (Rua and Alexandrescu 2024). To test for possible buffer effects on the titration we repeated the experiment using 5 mM HEPES buffer pH 7.0 and obtained a consistent K_d_ of (5.1 ± 0.7) x 10^-12^ M (Fig. S4).

The Co^2+^-binding experiment was done on an Ultrospec 8000 double-beam UV-Vis spectrophotometer (Thermo Fisher) using a 750 µL cuvette. The Z0 sample was 35 µM in 10 mM Tris pH 7.5, containing 0.5 mM TCEP. Data for Z0 were collected with and without 300 µM CoCl_2_. Each dataset was the average of 10 spectra, to improve the sensitivity in the 500-800 nm region of the spectrum where the weak d-d transition bands occur.

### 3.3 NMR spectroscopy and structure calculation

NMR assignments and structure determination were done using data collected on a 600 MHz Varian Innova spectrometer with a cryogenic probe. NMR samples had 40 µM Z0 containing 600 µM ZnSO_4_, dissolved in either 90% H2O/ 10% D_2_O (pH 6.3) or 99.98% D_2_O (pD 5.8). NMR samples had no other added buffers or salts. The temperature for all NMR experiments was maintained at 25 °C. For the Z0 sample dissolved in H_2_O we recorded 2D NOESY and TOCSY experiments with 200 ms and 70 ms mixing times, respectively. For the sample dissolved in D_2_O we recorded two NOESY experiments using mixing times of 200 and 50 ms, a 70 ms TOCSY experiment, and a ^1^H-^13^C HSQC experiment at natural abundance. NMR data analysis and assignments were performed with the program CcpNmr Analysis 2.5.2 (Vranken *et al*. 2005). The NOESY and TOCSY spectra enabled us to assign 91% of proton resonances, but because of the low sensitivity of the natural abundance ^1^H-^13^C HSQC experiment at 40 µM peptide concentrations we were only able to obtain assignments for ∼25% of the ^13^C NMR resonances expected from the amino acid sequence. ^1^H and ^13^C chemical shifts were referenced to an internal DSS (2,2-dimethyl-2-silapentane-5-sulfonate) standard.

For structure calculations, NOESY crosspeaks were uniformly assigned lower and upper distance bounds of 1.8 to 5.0 Å. Stereospecific methylene proton assignments were determined using a literature protocol (Case *et al*. 1994). Pseudoatom upper bound corrections of 1.0, 2.0, 2.4 and 1.5 Å were included for protons with ambiguous methylene, aromatic ring, prochiral methyl, and methyl group NMR signals, respectively (Wuthrich *et al*. 1983). Dihedral angle restraints were calculated from assigned chemical shifts using the program DANGLE (Cheung *et al*. 2010). Hydrogen bond restraints were included for amide protons in regular secondary structure based on the secondary structure consensus (Fig. 2B) and initial NMR structures excluding hydrogen bonds. The Zn^2+^ atom and ligands were restrained using distance bounds of 2.33-2.37 Å for Zn^2+^-Sγ and = 3.25-3.51 Å for Zn^2+^-Cβ for each of the three cysteines in the Z0 sequence, together with a 1.0-3.1 Å Zn^2+^-Nχ2 restraint for the sole histidine (Matousek and Alexandrescu 2004).

NMR structures calculations were performed on the NMRbox platform (Maciejewski *et al*. 2017) with the program X-Plor NIH v. 3.8, using the protein-4.0 parameter set (Bermejo *et al*. 2024). Structures were calculated using a distance geometry protocol followed by simulated annealing refinement using the X-plor script *prot_sa_refine_nogyr.inp* from the NESG site (https://nesgwiki.chem.buffalo.edu), as this gave less steric clashes for PSVS (Bhattacharya *et al*. 2007) and PDB validation. From an initial set of 38 structures with randomized phi and psi angles, the 20 lowest energy structures without violations were selected for deposition and analysis (Table S1).

### 3.4 ZNF sequence analysis

A PDB search was used to find ZNF domain fragments in complex with dsDNA for the *DNA-binders X-ray* dataset. To minimize the inclusion of redundant sequences from homologs of different species, we only included human proteins in this dataset. We used PDB-BLAST searches for each entry to exclude highly similar entries, such as multiple X-ray structures of the same protein. In cases where multiple PDB entries exist for different sized fragments of the same TF (e.g. PDB 5UND and 5K5H) we chose the structure encompassing the largest number of ZNF domains. Sequences were identified from the PDB entries and cross-checked with the matching UniProt entries to identify ZNF domain boundaries. The PDB and UniProt accession codes for each example, together with the ZNF domain boundaries and sequences to give a 28-a.a. alignment are listed in Table S2.

To construct the *DNA-binders Δ* and *Non-DNA Δ* datasets we did a PubMed search for papers describing TFs investigated through mutagenic deletion analysis to distinguish ZNFs involved and uninvolved in dsDNA binding. We identified 18 such publications contributing 57 DNA-binders and 91 non-binders. Because of the relatively small size of the literature database, we included human as well as sequences from other organisms but checked that these were non-redundant examples. For each TF we crosschecked the ZNF domain boundaries and sequences against the UniProt protein sequence database. The PubMed ID (PMID) codes for each publication, the UniProt accession IDs, UniProt ZNF boundaries, and sequences to obtain a 28-a.a. alignment are in Table S3.

To construct the *CCHC protein-binder* dataset we used a combination of literature papers, structures of CCHC ZNFs in the PDB, and CCHC-family ZNFs identified in the PROSITE database (Sigrist *et al*. 2013). To prevent redundancy all of the examples in the *CCHC protein-binder* dataset were human proteins. Altogether, we identified 28 CCHC-finger ZNFs from 18 different proteins. In previous work (Rua *et al*. 2023) we identified another 10 likely CCHC-finger examples that are currently annotated as degenerate by UniProt, but we did not include those examples because their folding status has unfortunately not yet been verified. The UniProt accession codes, domain boundaries and sequences for all the examples in the final CCHC dataset are given in Table S4.

Each of the ZNFs in the four datasets were scored with the ZF-prediction server (https://zf.princeton.edu/logoMain.php) (Persikov and Singh 2014). For the *CCHC* dataset a substantial number of examples (5/28), including the Z0 domain gave a ZF-server score of zero. Moreover, we noticed that CCHC-fingers intrinsically score 1-2 points lower in the program when the last Zn^2+^-ligand is a cysteine rather than a histidine. We therefore made a control dataset in which all the CCHC-fingers were artefactually changed to CCHH-fingers and all ZNFs scoring zero were removed. Even with this more stringent control dataset, the mean score (13.0 ± 5.5, n = 24) remains different from the other three groups at the p < 0.00001 significance level.

Sequence preferences for each dataset were investigated with sequence logos constructed from the individual ZNFs in each dataset. We aligned the sequences on the aromatic residue two positions before the first cysteine Zn^2+^-ligand (position 1), corresponding to the earliest conserved feature in the CCHH-finger ZNF motif (Klug 2010). With this alignment scheme, additional conserved aromatic and hydrophobic residues occur at positions 10 and 16, respectively, the four metal ligands have preferences for positions 3, 6, 19, and 23, and the ZNF residues in contact with nucleotide bases in dsDNA are at positions 12, 14, 15, and 18 as indicated in Fig. 3C. We set the upper boundary for the sequence motif at position 28 (Miller *et al*. 1985). In five examples of ZNFs that occur at the C-termini of proteins the sequence ends before position 28. To be able to use these sequences in our analysis we filled in the missing gaps with glycine residues.

### 3.5 Database accession numbers

NMR assignments and chemical shift values for the Z0 domain were deposited in the BMRB under accession code 52414. Coordinates and restraints for NMR structure calculations were deposited in the PDB under accession code 9BV0.

#### Note

During the writing of this research note, a paper describing the X-ray structure of a Z0 tetramer was published (Zheng *et al*. 2024) a week before the submission of our work to BioRxiv. No information from the tetramer structure (9B4P) was used for the NMR structure (9BV0), as our PDB and BMRB files were publicly released six months before the publication, and the crystal structure was still not available at the time of our deposition to BioRxiv. The monomer structures appear to be similar by NMR and crystallography, as both match the AlphaFold3 prediction for Z0. Besides small differences in peptide sequence and sample conditions, the tetramer was crystallized at a Z0 concentration (14 mg/ml) over 100-fold larger than that used in our NMR studies to prevent aggregation (0.13 mg/ml).

## Abbreviations

BCA: bicinchoninic acid
BCL11A: B-cell lymphoma/leukemia 11A
BMRB: biological magnetic resonance bank
CD: circular dichroism
dsDNA: double stranded deoxyribonucleic acid
EGTA: ethylene glycol-bis(β-aminoethyl ether)-*N,N,N*′,*N*′-tetraacetic acid
Hb: hemoglobin
HbA: adult hemoglobin
HbF: fetal hemoglobin
HEPES: 4-(2-hydroxyethyl)-1-piperazineethanesulfonic acid
HSQC: heteronuclear single quantum correlation
LMCT: ligand-to-metal charge transfer
NMR: nuclear magnetic resonance, NOE, nuclear Overhauser effect
NOESY: nuclear Overhauser effect spectroscopy
PDB: protein data bank
pLDDT: predicted local distance difference test
RMSD: root mean square deviation
SD: standard deviation
TCEP: tris(2-carboxyethyl)phosphine
TF: transcription factor
TOCSY: total correlation spectroscopy
UV-Vis: ultraviolet-visible spectrophotometry
Z0: CCHC zinc finger domain of BCL11A corresponding to residues 46-72 of UniProt entry Q9H165
ZNF: zinc finger.

## AUTHOR CONTRIBUTIONS

**R.E. Harris:** Investigation; data curation; formal analysis; visualization; writing – review and editing.

**R.D. Whitehead III:** Investigation; formal analysis; supervision; writing – review and editing;

**A.T. Alexandrescu:** Conceptualization; investigation; writing – original draft, review and editing; visualization; resources; project administration, supervision.

## ACKNOWLEDGEMENTS

We thank Prof. Carolyn Teschke for use of her group’s CD and UV-Vis spectrophotometers and Charley Damico for analysis of DNA-binding ZNF sequences during the early stages of this project.

## SUPPLEMENTARY DATA

Additional supporting information can be found online in the Supporting Information section at the end of this article.

