## Supporting_Information for "Solution structure of the Z0 domain from transcription repressor BCL11A sheds light on the sequence properties of protein-binding zinc-fingers"

**Rilee E. Harris, Richard D. Whitehead III, and Andrei T. Alexandrescu**

*Department of Molecular and Cellular Biology, University of Connecticut*

### Table of contents:

**Figure S1 – Temperature titration for $Zn^{2+}$-bound Z0.**
**Figure S2 – NMR spectra of $Zn^{2+}$-bound Z0 as a function of holo-peptide concentration.**
**Figure S3 – AlphaFold3 predictions of the Z0 dimer structure.**
**Figure S4 – Determination of $K_d$ for $Zn^{2+}$-binding in HEPES buffer.**

**Table S1 – Statistics for the 20 best NMR structures of the Z0 domain from BCL11A.**
**Table S2 – Dataset of dsDNA-binding ZNFs from X-ray structures.**
**Table S3 – Dataset of dsDNA-binding and non-binding ZNFs from literature deletion analyses.**
**Table S4 – Dataset of protein-binding CCHC ZNFs.**

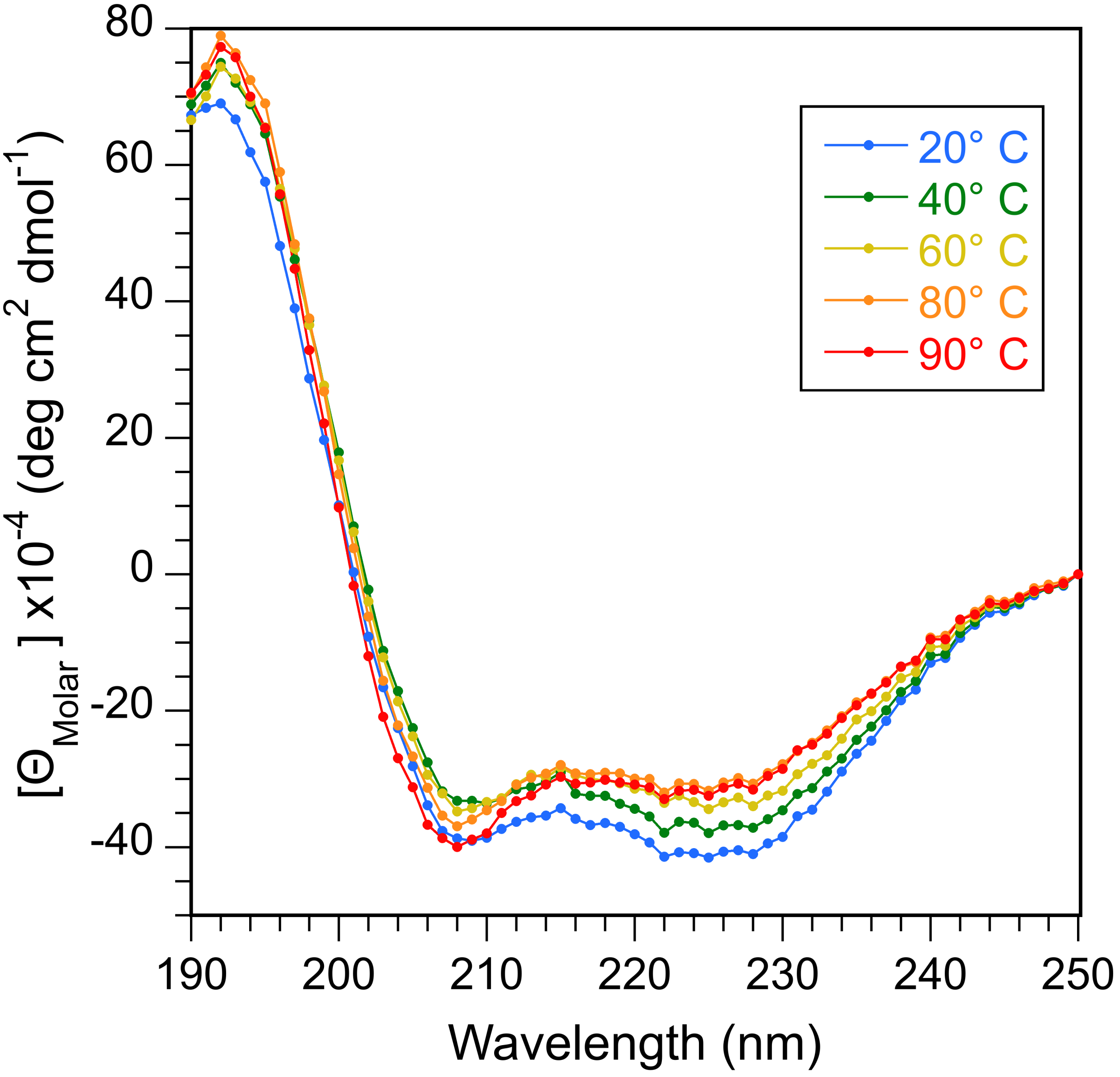


**Supplementary Figure S1 Temperature titration for $Zn^{2+}$-bound Z0.** Experiments were done with 40 μM Z0 in the presence of 50 μM $ZnSO_4$ at pH 7.0 containing 0.5 mM reducing agent TCEP.

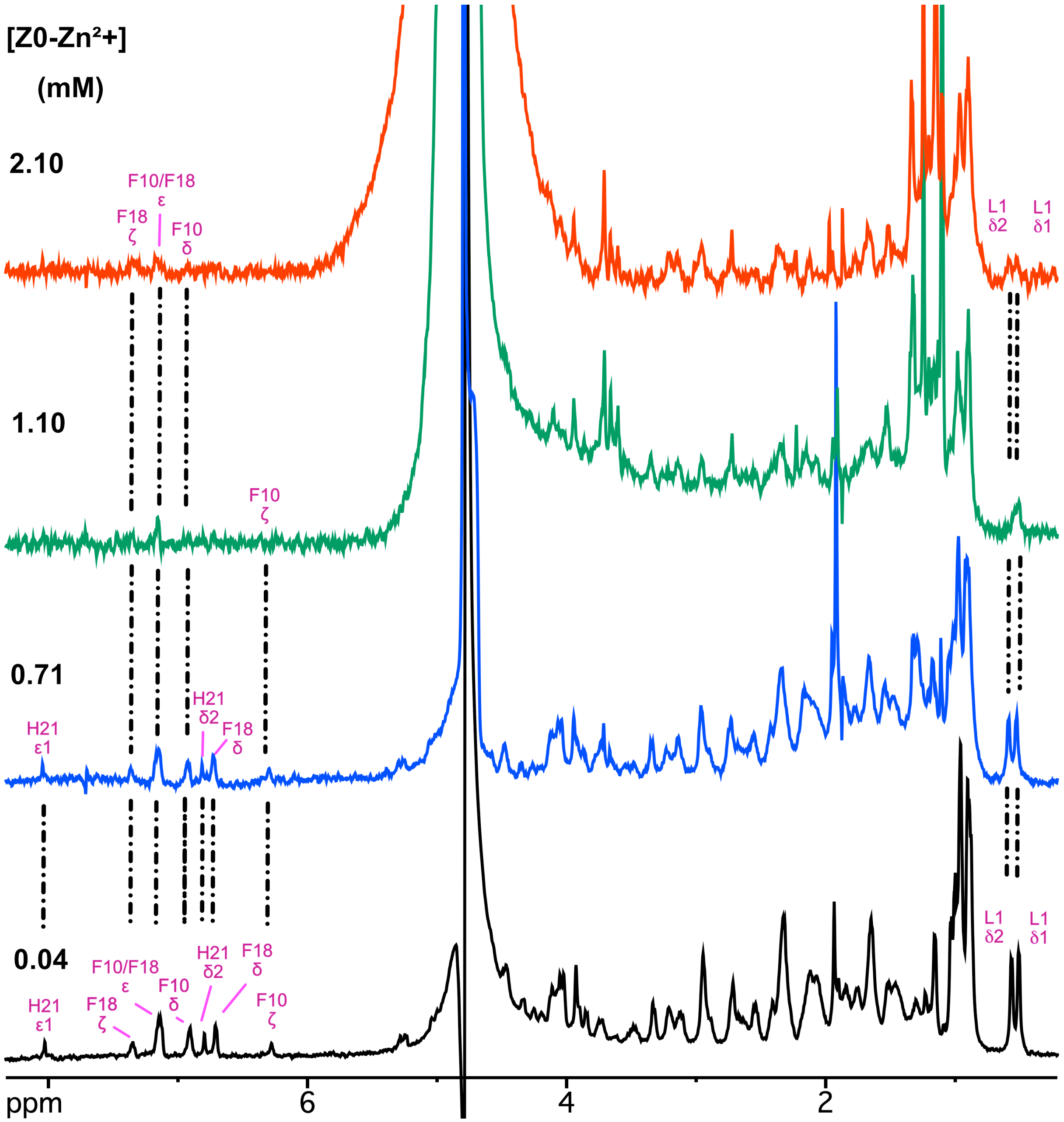


**Supplementary Figure S2 NMR spectra of $Zn^{2+}$-bound Z0 as a function of holo-peptide concentration.** A sample of 40 μM Z0 with equimolar $ZnSO_4$ was divided into aliquots and these were lyophilized so that when the peptide was re-dissolved in $D_2O$ the final concentrations were 0.04, 0.71, 1.10, and 2.10 mM. Resolved, assigned NMR signals from various parts of the protein were used to monitor the increase in NMR line width due to aggregation, however, broadening was non-selective for the subset of resolved resonances in the 1D spectra.

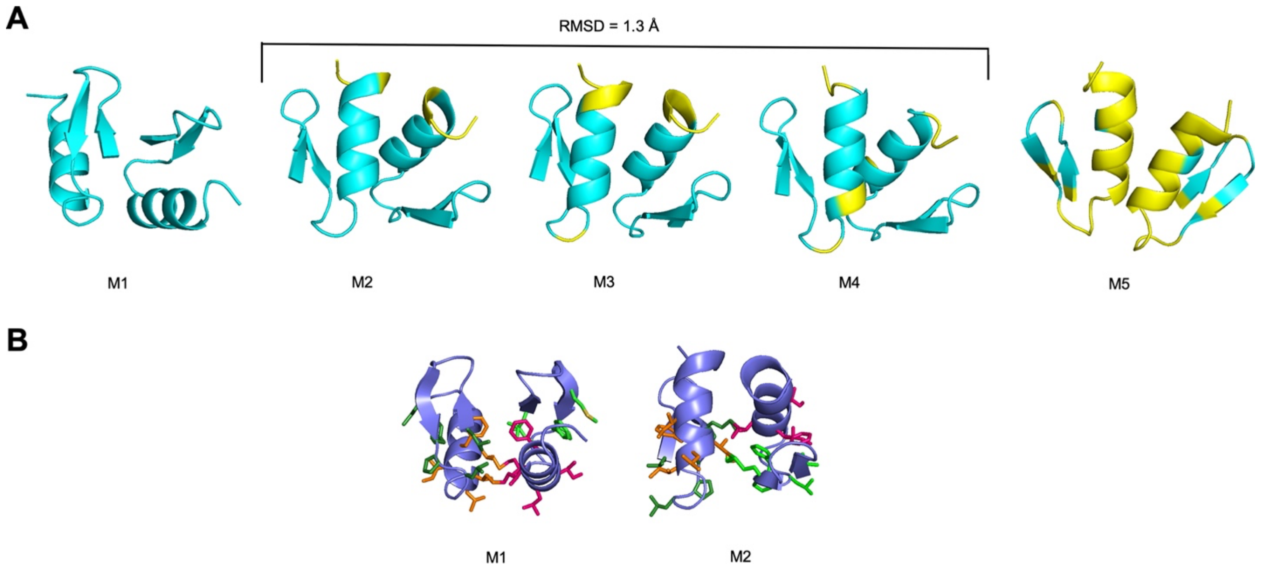


**Supplementary Figure S3  AlphaFold3 predictions of the Z0 dimer structure. (A)** The AlphaFold3 server (https://alphafoldserver.com) was used to predict Z0 dimer structures, which are colored by pLDDT score (yellow: low confidence 70> pLDDT >50, cyan: confident 90> pLDDT >70). Model M1 has the best scores rank of the five models but gives RMSDs of about 10 Å to the other models. Models M2-M3 form a structural cluster with an average RMSD of 1.3 Å amongst the three models. Model M5, which has the lowest confidence amongst the models, gives an average RMSD of 5Å to models M2-M4. (**B**) View of the predicted dimerization interface for models M1 and M2 showing hydrophobic residues L1, M8, 10-FPL-12 in light and dark green, and the hydrophobic region 15-ILIFI-19 at the N-terminus of the $\alpha$-helix in orange and red. In all five models the dimerization interface involves the highly hydrophobic N-terminus of the $\alpha$-helix. In model M1 this interface is more extensive than in the other models that have some of the hydrophobic residues positioned on the surface as illustrated for model M2.

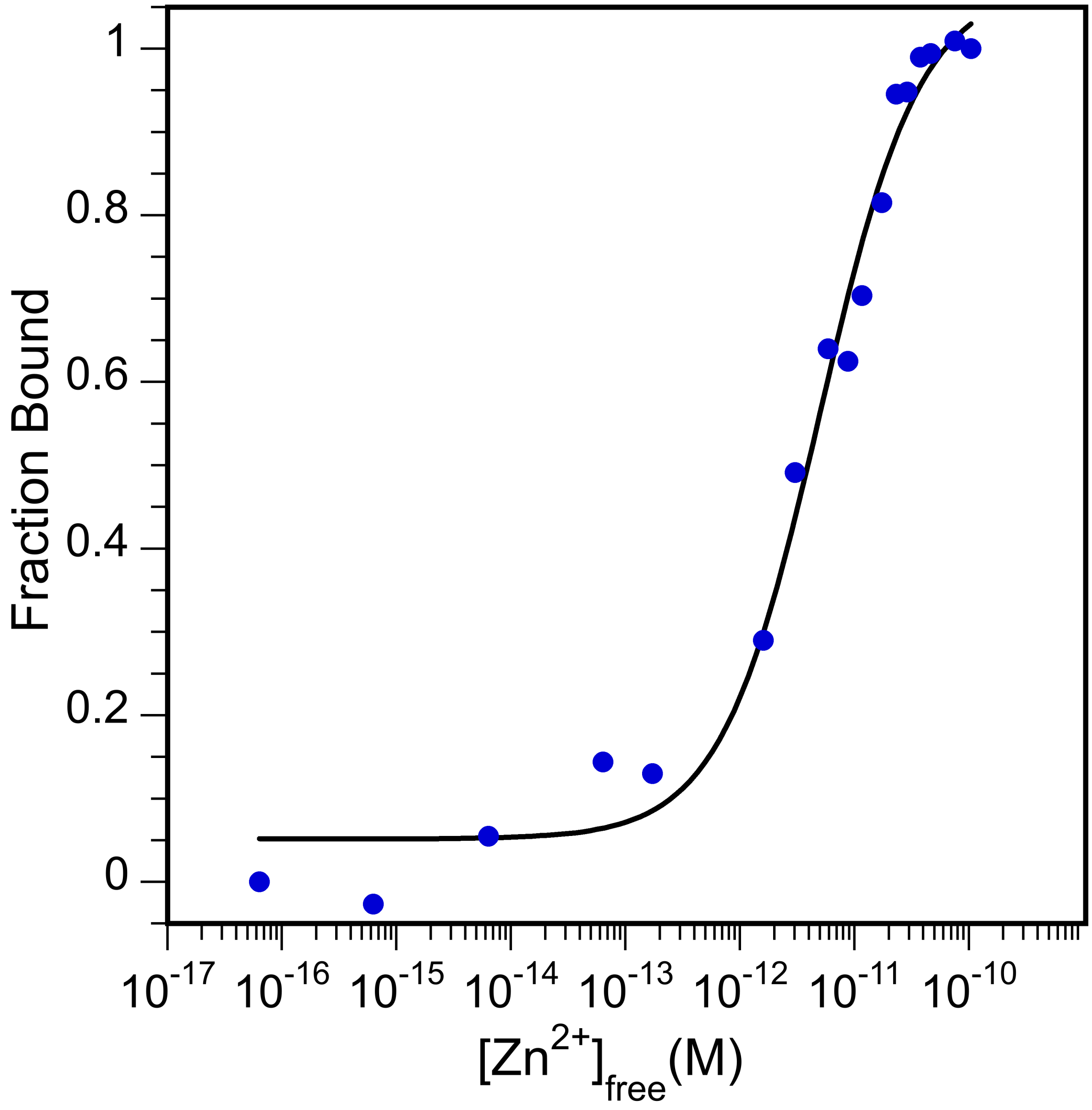


**Supplementary Figure S4 Determination of $K_d$ for $Zn^{2+}$-binding in HEPES buffer.** Experiments were done as described in the Methods except using 5 mM HEPES buffer pH 7.0 instead of 10 mM sodium phosphate buffer. The buffer also contained 0.2 mM TCEP and experiments were done at a temperature of 20 °C. For the titration, $ZnSO_4$ was added to a 40 μM Z0 sample containing a 10 mM concentration of the competitive chelator EGTA. The $K_d$ obtained in HEPES buffer was (5.1 ± 0.7) x $10^{-12}$ M.

**Table S1 Statistics for the 20 best NMR structures of the Z0 domain from BCL11A.**

| | | |
|---|---|---|
| NMR restraints (total) | 241 | |
| Distance (total) | 197 | |
| Intraresidue NOE (\|*i*-*j*\| = 0) | 69 | |
| Sequential NOE (\|*i*-*j*\| = 1) | 38 | |
| Medium-range NOE (1<\|*i*-*j*\|<5) | 32 | |
| Long-range NOE (\|*i*-*j*\|≥5) | 31 | |
| Hydrogen bond [a] | 20 | |
| Restraints to $Zn^{2+}$ atom [b] | 7 | |
| Dihedral ($\phi$ 17, $\psi$ 17, $\chi_1$ 10) | 44 | |
| *Restraint violations* | | |
| > 0.3 Å | 0 | |
| > 3° | 0 | |
| *Residual restraint violations* | | |
| Distance (Å) | 0.0338 ± 0.0032 [c] | |
| Dihedral (°) | 0.37 ± 0.10 | |
| *RMS deviations from ideal geometry* | | |
| Bonds (Å) | 0.00148 ± 0.00026 | |
| Angles (°) | 0.34 ± 0.04 | |
| Improper torsions (°) | 0.18 ± 0.02 | |
| *Ramachandran plot PROCHECK statistics for ordered residues* | | |
| Residues in most favored regions | 90.3 % | |
| Residues in allowed regions | 9.7 % | |
| Residues in generously allowed regions | 0.0 % | |
| Residues in disallowed regions | 0.0 % | |
| *Coordinate rms deviations (Å)* [d] | Backbone (Cα,N,C') | All |
| NMR ensemble to mean | 0.67 ± 0.13 | 1.50 ± 0.19 |
| NMR ensemble to Colabfold | 1.29 ± 0.18 | 2.06 ± 0.21 |
| NMR ensemble to PDB 8SXM 1-27 [e] | 2.06 ± 0.21 | N.A. |

[a]Two distance restraints (1.5-2.5 Å for NH-O and 2.5-3.5 Å for N-O) were included for each H-bond to enforce linearity.

[b]Two restraints per Cys (from Sγ and Cβ to maintain linearity) and one restraint per His (Nε2 rather than Nδ1, based on the histidine ring orientation in initial structures)

[c]All values are given as mean ± standard deviation.

[d]Alignments were done for the entire structure using the PyMol program with the align function and cycles=0. No atoms were rejected for the calculation of RMSD values.

[e]PDB 8SXM is the NMR structure of the CCHC zinc finger from ZNF750.

**Table S2 Dataset of dsDNA-binding ZNFs from X-ray structures.**

| UniProt ID | PDB ID | Uniprot a.a. | Sequence |
|---|---|---|---|
| Q9UJQ4 | 7Y31 | 870-892 | HGCTRCGKNFSSASALQIHERTHTGEKP |
| | | 898-920 | FVCNICGRAFTTKGNLKVHYMTHGANNN |
| Q9UJQ4 | 7Y3M | 382-404 | HKCKYCSKVFGTDSSLQIHLRSHTGERP |
| | | 410-432 | FVCSVCGHRFTTKGNLKVHFHRHPQVKA |
| P10074 | 5YJ3 | 521-544 | FSCEFCEQRFTEKGPLLRHVASRHQEGR |
| | | 550-572 | HFCQICGKTFKAVEQLRVHVRRHKGVRK |
| | | 578-600 | FECTECGYKFTRQAHLRRHMEIHDRVNY |
| Q96C55 | 7YSF | 114-136 | HFCPVCLRAFPYLSDLERHSISHSELKP |
| | | 142-164 | HQCKVCGKTFKRSSHLRRHCNIHAGLRP |
| | | 170-192 | FRCPLCPRRFREAGELAHHHRVHSGERP |
| | | 198-221 | YQCPICRLRFTEANTLRRHAKRKPEAMG |
| P49711 | 5UND | 351-373 | FKCSMCDYASVEVSKLKRHIRSHTGERP |
| | | 379-401 | FQCSLCSYASRDTYKLKRHMRTHSGEKP |
| | | 407-430 | YECYICHARFTQSGTMKMHILQKTENVA |
| | | 437-460 | FHCPHCDTVIARKSDLGVHLRKQSYIEQ |
| | | 467-489 | KKCRYCDAVFHERYALIQHQKSHKNEKR |
| | | 495-517 | FKCDQCDYACRQERHMIMHKRTHTGEKP |
| | | 523-546 | YACSHCDKTFRQKQLLDMHFKRYDPNFV |
| P49711 | 5K5H | 351-373 | FKCSMCDYASVEVSKLKRHIRSHTGERP |
| | | 379-401 | FQCSLCSYASRDTYKLKRHMRTHGESKP |
| | | 407-430 | YECYICHARFTQSGTMKMHILQKTENVA |
| P49711 | 5K5I | 379-401 | FQCSLCSYASRDTYKLKRHMRTHSGEKP |
| | | 407-430 | YECYICHARFTQSGTMKMHILQKTENVA |
| | | 437-460 | FHCPHCDTVIARKSDLGVHLRKQSYIEQ |
| | | 467-489 | KKCRYCDAVFHERYALIQHQKSHKNEKR |
| P49711 | 5K5L | 407-430 | YECYICHARFTQSGTMKMHILQKTENVA |
| | | 437-460 | FHCPHCDTVIARKSDLGVHLRKQSYIEQ |
| | | 467-489 | KKCRYCDAVFHERYALIQHQKSHKNEKR |
| P49711 | 5KKQ | 322-345 | HKCPDCDMAFVTSGELVRHRRYKTHEKP |
| | | 351-373 | FKCSMCDYASVEVSKLKRHIRSHTGERP |
| | | 379-401 | FQCSLCSYASRDTYKLKRHMRTHSGEKP |

| | | | |
|---|---|---|---|
| | | 407-430 | YECYICHARFTQSGTMKMHILQKTENVA |
| | | 437-460 | FHCPHCDTVIARKSDLGVHLRKQSYIEQ |
| P49711 | 5T0U | 294-316 | HKCHLCGRAFRTVTLLRNHLNTHLSEVN |
| | | 322-345 | HKCPDCDMAFVTSGELVRHRRYKTHEKP |
| | | 351-373 | FKCSMCDYASVEVSKLKRHIRSHTGERP |
| | | 379-401 | FQCSLCSYASRDTYKLKRHMRTHSGEKP |
| | | 407-430 | YECYICHARFTQSGTMKMHILQKTENVA |
| | | 437-460 | FHCPHCDTVIARKSDLGVHLRKQSYIEQ |
| P49711 | 5YEF | 294-316 | HKCHLCGRAFRTVTLLRNHLNTHTGTRP |
| | | 322-345 | HKCPDCDMAFVTSGELVRHRRYKTHEKP |
| | | 351-373 | FKCSMCDYASVEVSKLKRHIRSHTGERP |
| | | 379-401 | FQCSLCSYASRDTYKLKRHMRTHSGEKP |
| | | 407-430 | YECYICHARFTQSGTMKMHILQKTENVA |
| | | 437-460 | FHCPHCDTVIARKSDLGVHLRKQSYIEQ |
| | | 467-489 | KKCRYCDAVFHERYALIQHQKSHKNEKR |
| P49711 | 5YEH | 351-373 | FKCSMCDYASVEVSKLKRHIRSHTGERP |
| | | 379-401 | FQCSLCSYASRDTYKLKRHMRTHSGEKP |
| | | 407-430 | YECYICHARFTQSGTMKMHILQKTENVA |
| | | 437-460 | FHCPHCDTVIARKSDLGVHLRKQSYIEQ |
| | | 467-489 | KKCRYCDAVFHERYALIQHQKSHKNEKR |
| P49711 | 5YEL | 407-430 | YECYICHARFTQSGTMKMHILQKTENVA |
| | | 437-460 | FHCPHCDTVIARKSDLGVHLRKQSYIEQ |
| | | 467-489 | KKCRYCDAVFHERYALIQHQKSHKNEKR |
| | | 495-517 | FKCDQCDYACRQERHMIMHKRTHTGEKP |
| | | 523-546 | YACSHCDKTFRQKQLLDMHFKRYDPNFV |
| | | 555-577 | FVCSKCGKTFTRRNTMARHADNCAGPDG |
| Q86VK4 | 6WMI | 219-243 | LKCTVEGCDRTFVWPAHFKYHLKRNDRS |
| | | 249-273 | FICPAEGCGKSFYVLQRLKVHMRNGEKP |
| | | 279-303 | FMCHESGCGKQFTTAGNLKNHRRTGEKP |
| | | 309-333 | FLCEAQGCGRSFAEYSSLRKHLVSGEKP |
| | | 339-362 | HQCQVCGKTFSQSGSRNVHMRKHLQLGA |
| P18146 | 4R2A | 338-362 | YACPVESCDRRFSRSDELTRHIRTGQKP |
| | | 368-390 | FQCRICMRNFSRSDHLTTHIRTHTGEKP |
| | | 396-418 | FACDICGRKFARSDERKRHTKIHLRQKD |
| O95365 | 7EYI | 382-404 | QKCPICEKVIQGAGKLPRHIRTHTGEKP |

| | | | |
|---|---|---|---|
| | | 410-432 | YECNICKVRFTRQDKLKVHMRKHTGEKP |
| | | 438-460 | YLCQQCGAAFAHNYDLKNHMRVHTGLRP |
| | | 466-490 | YQCDSCCKTFVRSDHLHRHLKKDNGVPS |
| O95365 | 7N5S | 382-404 | QKCPICEKVIQGAGKLPRHIRTHTGEKP |
| | | 410-432 | YECNICKVRFTRQDKLKVHMRKHTGEKP |
| | | 438-460 | YLCQQCGAAFAHNYDLKNHMRVHTGLRP |
| | | 466-490 | YQCDSCCKTFVRSDHLHRHLKKDNGVPS |
| Q9H165 | 6U9Q | 742-764 | DTCEYCGKVFKNCSNLTVHRRSHTGERP |
| | | 770-792 | YKCELCNYACAQSSKLTRHMKTHGQVGK |
| | | 800-823 | YKCEICKMPFSVYSTLEKHMKKWSDRVL |
| Q86T24 | 4F6M | 494-516 | YICIVCKRSYVCLTSLRRHFNIHSWEKK |
| | | 522-544 | YPCRYCEKVFPLAEYRTKHEIHHTGERR |
| | | 550-573 | YQCLACGKSFINYQFMSSHIKSVSQDPS |
| Q96DT7 | 8GN3 | 722-744 | LKCPHCSYVAKYRRTLKRHLLIHTGVRS |
| | | 750-772 | FSCDICGKLFTRREHVKRHSLVHKKDKK |
| O43474 | 6VTX | 430-454 | HTCDYAGCGKTYTKSSHLKAHLRTGEKP |
| | | 460-484 | YHCDWDGCGWKFARSDELTRHYRTGHRP |
| | | 490-512 | FQCQKCDRAFSRSDHLALHMKRHF |
| O75362 | 3UK3 | 471-493 | RECSYCGKFFRSNYYLNIHLRTHTGEKP |
| | | 499-521 | YKCEFCEYAAAQKTSLRYHLERHHKEKQ |
| P25490 | 1UBD | 296-320 | IACPHKGCTKMFRDNSAMRKHLHGPRVH |
| | | 325-347 | HVCAECGKAFVESSKLKRHQLVHTGEKP |
| | | 353-377 | FQCTFEGCGKRFSLDFNLRTHVRTGDRP |
| | | 383-407 | YVCPFDGCNKKFAQSTNLKSHILAKAKN |
| Q8NAP3 | 6 E93 | 1038-1060 | YQCKTCGRCFSVQGNLQKHERIHLGLKE |
| | | 1066-1088 | FVCQYCNKAFTLNETLKIHERIHTGEKR |
| | | 1094-1116 | YHCQFCFQRFLYLSTKRNHEQRHIREHN |
| Q9NQV7 | 5EGB | 720-742 | YVCRECGRGFSNKSHLLRHQRTHTGEKP |
| | | 748-770 | YVCRECGRGFRDKSHLLRHQRTHTGEKP |
| | | 776-798 | YVCRECGRGFRDKSNLLSHQRTHTGEKP |
| | | 804-826 | YVCRECGRGFSNKSHLLRHQRTHTGEKP |
| | | 832-854 | YVCRECGRGFRNKSHLLRHQRTHTGEKP |

| | | | |
|---|---|---|---|
| Q96JB3 | 7TXC | 505-532 | FKCSVCEKTYKDPATLRQHEKTHFPCNI |
| | | 533-560 | FPCNICGKMFTQRGTMTRHMRSHFACDE |
| | | 561-588 | FACDECGMRFTRQYRLTEHMRVHYECQL |
| | | 589-615 | YECQLCGGKFTQQRNLISHLRMH |

**Table S3 Dataset of dsDNA-binding and non-binding ZNFs from literature deletion analyses.**

| PMID | Uniprot ID | a.a. # | DNA-binding ZNF sequences | a.a. # | non-DNA-binding ZNF sequences |
|---|---|---|---|---|---|
| 29429977 | P17010 | 719-741 | FRCKRCRKGFRQQSELKKHMKTHSGRKV | 425-447 | YPCMICGKKFKSRGFLKRHMKNHPEHLA |
| | | 747-770 | YQCEYCEYSTTDASGFKRHVISIHTKDY | 456-478 | YRCTDCDYTTNKKISLHNHLESHKLTSK |
| | | 776-798 | HRCEYCKKGFRRPSEKNQHIMRHHKEVG | 488-510 | IECDECGKHFSHAGALFTHKMVHKEKGA |
| | | | | 519-542 | HKCKFCEYETAEQGLLNRHLLAVHSKNF |
| | | | | 548-570 | HICVECGKGFRHPSELKKHMRIHTGEKP |
| | | | | 576-599 | YQCQYCEYRSADSSNLKTHVKTKHSKEM |
| | | | | 605-627 | FKCDICLLTFSDTKEVQQHALIHQESKT |
| | | | | 633-656 | HQCLHCDHKSSNSSDLKRHIISVHTKDY |
| | | | | 662-684 | HKCDMCDKGFHRPSELKKHVAAHKGKKM |
| | | | | 690-713 | HQCRHCDFKIADPFVLSRHILSVHTKDL |
| 28500257 | P10074 | 578-600 | FECTECGYKFTRQAHLRRHMEIHDRVEN | 291-313 | VECPTCHKKFLSKYYLKVHNRKHTGEKP |
| | | | | 319-344 | FECPKCGKCYFRKENLLEHEARNCMNRS |
| | | | | 350-372 | FTCSVCQETFRRRMELRVHMVSHTGEMP |
| | | | | 378-401 | YKCSSCSQQFMQKKDLQSHMIKLHGAPK |
| | | | | 407-430 | HACPTCAKCFLSRTELQLHEAFKHRGEK |
| | | | | 436-459 | FVCEECGHRASSRNGLQMHIKAKHRNER |
| | | | | 465-487 | HVCEFCSHAFTQKANLNMHLRTHTGEKP |
| | | | | 493-515 | FQCHLCGKTFRTQASLDKHNRTHTGERP |
| | | | | 521-544 | FSCEFCEQRFTEKGPLLRHVASRHQEGR |
| | | | | 550-572 | HFCQICGKTFKAVEQLRVHVRRHKGVRK |
| 7604032 | P13360 | 493-515 | FRCPICDRRFSQSSSVTTHMRTHSGERP | 437-459 | NLCRLCGKTYARPSTLKTHLRTHSGERP |
| | | 521-542 | YRCSSCKKSFSDSSTLTKHLRIHHSGEK | 465-487 | YRCPDCNKSFSQAANLTAHVRTHTGQKP |
| | | 549-571 | YQCKLCLLRFSQSGNLNRHMRVHGNNNS | | |
| 8378770 | P08151 | 268-295 | FVCHWGGCSRELRPFKAQYMLVVHMRRH | 235-260 | TDCRWDGCSQEFDSQEQLVHHINSEHIH |
| | | 301-325 | HKCTFEGCRKSYSRLENLKTHLRSHTGE | | |
| | | 331-356 | YMCEHEGCSKAFSNASDRAKHQNRTHSN | | |
| 1569931 | O75626 | 575-597 | YECNVCAKTFGQLSNLKVHLRVHSGERP | 631-653 | HECQVCHKRFSSTSNLKTHLRLHSGEKP |
| | | 603-625 | FKCQTCNKGFTQLAHLQKHYLVHTGEKP | 659-681 | YQCKVCPAKFTQFVHLKLHKRLHTRERP |
| 19853363 | P19544 | 353-377 | YQCDFKDCERRFSRSDQLKRHQRRHTGV | 323-347 | FMCAYPGCNKRYFKLSHLQMHSRKHTGE |
| | | 383-405 | FQCKTCQRKFSRSDHLKTHTRTHTGKTS | 414-438 | FSCRWPSCQKKFARSDELVRHHNMHQRN |
| 8662611 | P39933 | 49-74 | YFCDYDGCDKAFTRPSILTEHQLSVHQG | 194-219 | YQCTFAGCCKEFRIWSQLQSHIKNDHPK |
| | | 80-102 | FQCDKCAKSFVKKSHLERHLYTHSDTKP | 222-244 | LKCPICSKPCVGENGLQMHMIIHDDSLV |
| | | 108-130 | FQCSYCGKGVTTRQQLKRHEVTHTKSFI | 253-277 | WKCHICPDMSFSRKHDLLTHYGSIHTEE |
| | | 134-159 | FICPEEGCNLRFYKHPQLRAHILSVHLH | 365-389 | YRCFYNNCSRTFKTKEKYEKHIDKHKVH |

| | | | | | |
|---|---|---|---|---|---|
| | | 163-186 | LTCPHCNKSFQRPYRLRNHISKHHDPEV | | |
| 9578568 | P08047 | 656-680 | FMCTWSYCGKRFTRSDELQRHKRTHTGE | 626-650 | HICHIQGCGKVYGKTSHLRAHLRWHTGE |
| | | 686-708 | FACPECPKRFMRSDHLSKHIKTHQNKKG | | |
| 29606353 | Q9H165 | 770-792 | YKCELCNYACAQSSKLTRHMKTHGQVGK | 170-193 | YTCTTCKQPFTSAWFLLQHAQNTHGLRI |
| | | 800-823 | YKCEICKMPFSVYSTLEKHMKKWHSDRV | 377-399 | KSCEFCGKTFKFQSNLVVHRRSHTGEKP |
| | | | | 405-429 | YKCNLCDHACTQASKLKRHMKTHMHKSS |
| | | | | 742-764 | DTCEYCGKVFKNCSNLTVHRRSHTGERP |
| 17827499 | P49711 | 351-373 | FKCSMCDYASVEVSKLKRHIRSHTGERP | 266-288 | FQCELCSYTCPRRSNLDRHMKSHTDERP |
| | | 379-401 | FQCSLCSYASRDTYKLKRHMRTHSGEKP | 294-316 | HKCHLCGRAFRTVTLLRNHLNTHTGTRP |
| | | 407-430 | YECYICHARFTQSGTMKMHILQKHTENV | 322-345 | HKCPDCDMAFVTSGELVRHRRYKHTHEK |
| | | 437-460 | FHCPHCDTVIARKSDLGVHLRKQHSYIE | 467-489 | KKCRYCDAVFHERYALIQHQKSHKNEKR |
| | | | | 495-517 | FKCDQCDYACRQERHMIMHKRTHTGEKP |
| | | | | 523-546 | YACSHCDKTFRQKQLLDMHFKRYHDPNF |
| | | | | 555-577 | FVCSKCGKTFTRRNTMARHADNCAGPDG |
| 11145971 | Q13127 | 332-355 | FKCDQCSYVASNQHEVTRHARQVHNGPK | 159-181 | FRCKPCQYEAESEEQFVHHIRVHSAKKF |
| | | 361-383 | LNCPHCDYKTADRSNFKKHVELHVMPRQ | 216-238 | IRCDRCGYNTNRYDHYTAHLKHHTRAGD |
| | | 389-412 | FNCPVCDYAASKKCNLQYHFKSKHPTCP | 248-270 | YKCIICTYTTVSEYHWRKHLRNHFPRKV |
| | | | | 276-298 | YTCGKCNYFSDRKNNYVQHVRTHTGERP |
| | | | | 304-326 | YKCELCPYSSSQKTHLTRHMRTHSGEKP |
| | | | | 1060-1082 | FVCIFCDRSFRKGKDYSKHLNRHLVNVY |
| 11154279 | A0JPL0 | 295-317 | FQCPYCGNSFRRKSYLIEHQRIHTGEKP | 211-233 | FECDECDSSFLMTEVAFPHDRAHRGVRD |
| | | 323-345 | YICSQCGKAFRQKTALTLHEKTHTDGKP | | |
| | | 351-373 | YLCVDCGKSFRQKATLTRHHKTHTGEKA | | |
| | | 379-401 | YECTQCGSAFGKKSYLIDHQRTHTGEKP | | |
| | | 407-429 | YQCAECGKAFIQKTTLTVHQRTHTGEKP | | |
| | | 435-457 | YMCSECGKSFCQKTTLTLHQRIHTGEKP | | |
| | | 463-485 | YVCSDCGKSFRQKAILTVHYRIHTGEKS | | |
| | | 491-513 | NGCPQCGKAFSRKSNLIRHQKTHTGEKP | | |
| 11056018 | Q14872 | 140-164 | YQCTFEGCPRTYSTAGNLRTHQKTHRGE | 259-283 | FRCDHDGCGKAFAASHHLKTHVRTHTGE |
| | | 170-194 | FVCNQEGCGKAFLTSYSLRIHVRVHTKE | 289-313 | FFCPSNGCEKTFSTQYSLKSHMKGHDNK |
| | | 200-224 | FECDVQGCEKAFNTLYRLKAHQRLHTGK | | |
| | | 229-253 | FNCESEGCSKYFTTLSDLRKHIRTHTGE | | |
| 21765982 | Q13422 | 145-167 | FQCNQCGASFTQKGNLLRHIKLHSGEKP | 117-139 | LKCDICGIICIGPNVLMVHKRSHTGERP |
| | | 173-195 | FKCHLCNYACRRRDALTGHLRTHSVGKP | 201-224 | HKCGYCGRSYKQRSSLEEHKERCHNYLE |
| | | | | 462-484 | YKCEHCRVLFLDHVMYTIHMGCHGFRDP |

| | | | | | |
|---|---|---|---|---|---|
| | | | | 490-514 | FECNMCGYHSQDRYEFSSHITRGEHRFH |
| 8754800 | Q99684 | 312-334 | FDCKICGKSFKRSSTLSTHLLIHSDTRP | 255-278 | YKCIKCSKVFSTPHGLEVHVRRSHSGTR |
| | | 340-362 | YPCQYCGKRFHQKSDMKKHTFIHTGEKP | 284-306 | FACEMCGKTFGHAVSLEQHKAVHSQERS |
| | | 368-390 | HKCQVCGKAFSQSSNLITHSRKHTGFKP | 396-419 | FGCDLCGKGFQRKVDLRRHRETQHGLK |
| 12549926 | P47043 | 768-790 | YKCKTCKRCFSSEETLVQHTRTHSGEKP | 579-604 | LKCKWKECPESCSSLFDLQRHLLKDHVS |
| | | 796-818 | YKCHICNKKFAISSSLKIHIRTHTGEKP | 616-641 | LACNWEDCDFLGDDTCSIVNHINCQHGI |
| | | | | 705-730 | VICQWDGCNKSFSSAQELNDHLEAVHLT |
| | | | | 738-762 | YQCLWHDCHRTFPQRQKLIRHLKVHSKY |
| | | | | 824-846 | LQCKICGKRFNESSNLSKHIKTHTHQKK |
| 10660046 | Q2M1K9 | 138-160 | YPCQFCDKSFIRLSYLKRHEQIHSDKLP | 67-93 | YTCDHCQQDFESLADLTDHRAHRCPGDG |
| | | 166-188 | FKCTYCSRLFKHKRSRDRHIKLHTGDKK | 409-433 | YSCPYCSKRDFNSLAVLEIHLKTIHADK |
| | | 194-216 | YHCHECEAAFSRSDHLKIHLKTHSSSKP | 441-464 | HTCQICLDSMPTLYNLNEHVRKLHKNHA |
| | | 222-244 | FKCTVCKRGFSSTSSLQSHMQAHKKNKE | 480-503 | FHCNYCPEMFADINSLQEHIRVSHCGPN |
| | | 263-286 | FMCDYCEDTFSQTEELEKHVLTRHPQLS | 517-540 | FFCNQCSMGFLTESSLTEHIQQAHCSVG |
| | | 295-318 | LQCIHCPEVFVDENTLLAHIHQAHANQK | 563-588 | YSCPYCTNSPIFGSILKLTKHIKENHKN |
| | | 323-345 | HKCPMCPEQFSSVEGVYCHLDSHRQPDS | 632-654 | YPCNQCDLKFSNFESFQTHLKLHLELLL |
| | | | | 662-684 | QACPQCKEDFDSQESLLQHLTVHYMTTS |
| | | | | 692-715 | YVCESCDKQFSSVDDLQKHLLDMHTFVL |
| | | | | 720-743 | YHCTLCQEVFDSKVSIQVHLAVKHSNEK |
| | | | | 750-773 | YRCTACNWDFRKEADLQVHVKHSHLGNP |
| | | | | 781-803 | HKCIFCGETFSTEVELQCHITTHSKKYN |
| | | | | 807-830 | YNCKFCSKAFHAIILLEKHLREKHCVFD |
| | | | | 886-908 | YGCDICGAAYTMEVLLQNHRLRDHNIRP |
| | | | | 930-952 | HKCNVCSRTFFSENGLREHLQTHRGPAK |
| | | | | 959-981 | YMCPICGERFPSLLTLTEHKVTHSKSLD |
| | | | | 1020-1042 | FRCVVCMQTVTSTLELKIHGTFHMQKLA |
| | | | | 1064-1082 | YKCALCLKEFRSKQDLVKLDVNGLPYGL |
| | | | | 1120-1143 | LRCPECSVKFESAEDLESHMQVDHRDLT |
| | | | | 1168-1190 | YQCIKCQMTFENEREIQIHVANHMIEEG |
| | | | | 1198-1220 | HECKLCNQMFDSPAKLLCHLIEHSFEGM |
| | | | | 1229-1252 | FKCPVCFTVFVQANKLQQHIFAVHGQED |
| | | | | 1259-1282 | YDCSQCPQKFFFQTELQNHTMSQHAQ |
| 23059534 | Q9NU63 | 147-169 | FCCTLCDKTYCDASGLSRHRRVHLGYRP | 91-113 | FFCYTCGKCFSRRSYLYSHQFVHNPKLT |
| | | 175-197 | HSCSVCGKSFRDQSELKRHQKIHQNQEP | 119-141 | NSCSQCGKLFRSPKSLSYHRRMHLGERP |
| | | | | 300-322 | FCCPHCSLTFSKKSYLSRHQKAHLTEPP |
| | | | | 328-350 | NYCFHCSKSFSSFSRLVRHQQTHWKQKS |
| | | | | 356-378 | YLCPICDLSFGEKEGLMDHWRGYKGKDL |

**Table S4 Dataset of protein-binding CCHC ZNFs.**

| UniProt ID | Uniprot a.a. | Sequence |
|---|---|---|
| Q8IX07 | 235 - 268 | FPCKDCGIWYRSERNLQAHLLYYCASRQ |
| | 571 - 604 | ATCFECEITFSNVNNYYVHKRLYCSGRR |
| | 677 - 710 | TLCEACNIRFSRHETYTVHKRYYCASRH |
| | 811 - 844 | HECTACRVSFHSLEAYLAHKKYSCPAAP |
| | 968 - 1001 | RYCRLCNIKFSSLSTFIAHKKYYCSSHA |
| Q8WW38 | 244 - 277 | FPCKSCGIWYRSERNLQAHLMYYCSGRQ |
| | 542 - 575 | ATCFECNITFNNLDNYLVHKKHYCSSRW |
| | 681 - 714 | TTCEACNITFSRHETYMVHKQYYCATRH |
| | 848 - 881 | HECTVCKISFNKVENYLAHKQNFCPVTA |
| | 1113 - 1146 | KYCRLCDIQFNNLSNFITHKKFYCSSHA |
| P15822 | 956 - 986 | FECETCRNRYRKLENFENHKKFYCSELH |
| Q5T1R4 | 640 - 670 | YECNICGARYKKRDNYEAHKKYYCSELQ |
| Q9Y6K9 | 389 - 419 | FCCPKCQYQAPDMDTLQIHVMECIE |
| Q96CV9 | 547 - 577 | HSCPKCGEVLPDIDTLQIHVMDCII |
| Q8NFZ5 | 397 - 429 | LQCPHCLQCFSDEQGEELLRHVAECCQ |
| Q6NZ36 | 144 - 180 | RSCPMCQKEFAPRLTQLDVDSHLAQCLA |
| Q6PJP8 | 119 - 149 | GYCPNCQMPFSSLIGQTPRWHVFECLDS |
| Q9Y2M0 | 41 - 69 | LACPVCSKMVPRYDLNRHLDEMCANNDF |
| Q9UBT6 | 621 - 651 | LTCPVCFRAQGCISLEALNKHVDECLDG |
| | 776 - 806 | LVCPVCNVEQKTSDLTLFNVHVDVCLNK |
| Q9NS91 | 201 - 228 | VDCPVCGVNIPESHINKHLDSCLSREEK |
| Q8IY92 | 293 - 323 | FFCQICQKNLSAMNVTRREQHVNRCLDE |
| | 333 - 361 | PECPICGKPFLTLKSRTSHLKQCAVKME |

| | | |
|---|---|---|
| Q9H040 | 453 - 480 | VNCPVCQNEVLESQINEHLDWCLEGDSI |
| Q96RL1 | 502 - 529 | VSCPLCDQCFPPTKIERHAMYCNGLMEE |
| Q96S55 | 17 - 44 | VQCPVCQQMMPAAHINSHLDRCLLLHPA |
| Q32MQ0 | 25 - 51 | FQCPFTCNEKSHLFNHMKYGLCKNSITL |
| Q13064 | 266 - 293 | CDMCGLQTLHPMDAAQREEHMRACIEAH |
